# Distinct roles of neuronal phenotypes during neurofeedback adaptation

**DOI:** 10.1101/2025.05.02.651867

**Authors:** Yi Zhao, Hannah M. Stealey, Hung-Yun Lu, Enrique Contreras-Hernandez, Yin-Jui Chang, Philippe Tobler, Samantha R. Santacruz

**Affiliations:** Department of Biomedical Engineering, University of Texas at Austin; Austin, 78712, Texas, USA; Department of Economics, University of Zurich; Zurich CH-8006, Switzerland; Neuroscience Center Zurich, University of Zurich and Swiss Federal Institute of Technology Zurich; Zurich CH-8006, Switzerland; Department of Electrical and Computer Engineering, University of Texas at Austin; Austin, 78712, Texas, USA; Interdisciplinary Neuroscience Program, University of Texas at Austin; Austin, 78712, Texas, USA

## Abstract

Learning adaptation allows the brain to refine motor patterns in response to changing environments rapidly. While population-level neural dynamics and single-neuron activity in motor learning have been widely studied, the contributions of individual neuron types remain poorly understood. Here, we employed a brain-machine interface (BMI) task with perturbations of varying difficulty to investigate single-neuron dynamics underlying neurofeedback adaptation in two rhesus macaques. Cortical neurons were classified based on waveform shape into narrow waveform (NW) and broad waveform (BW) categories, representing putative inhibitory interneurons and excitatory pyramidal neurons, respectively. Compared to BW neurons, NW neurons were more active and more strongly involved in the learning process. Moreover, task difficulty modulated neural responsiveness and coordination within both neuron groups, highlighting differential neuron engagement during motor learning. Our findings provide novel insights into single-neuron mechanisms underlying neurofeedback adaptation and emphasize the distinct functional roles of neuronal phenotypes in rapid learning processes.

**Author Summary:** Understanding how the brain adapts to changes is crucial for improving treatments for neurological disorders and enhancing brain-machine interface (BMI) technology, which allows control of external devices using neural signals. In our study, we investigated how different types of brain cells contribute to the adaptation process when individuals encounter unexpected challenges during neurofeedback tasks. We discovered that one type of neuron was notably more active and played a more engaged role in rapidly adjusting neural activity compared to another type. These neurons demonstrated stronger coordination in their activity and showed greater responsiveness as the task difficulty increased. Our findings highlight the distinct roles of specific neurons in quickly adapting to neurofeedback tasks, offering insights that could enhance therapies for movement disorders and improve the precision and reliability of brain-controlled prosthetics.

## Introduction

The ability to adapt motor skills to new situations exemplifies the brain’s capacity for motor learning adaptation, a process that enables humans and animals to adjust and refine motor patterns in response to changing environments or physical capabilities. For example, after learning to play tennis, adapting to a mild wind while hitting the ball is more manageable than adjusting to strong wind conditions because the brain can quickly adapt to smaller perturbations. Motor learning adaptation relies on error-driven learning mechanisms that modify neural patterns in motor areas of the brain, facilitating rapid adjustments to external changes[1–5]. Researchers have studied neural mechanisms, such as neuroplasticity, to provide insights into how the brain learns and modifies its structure and function during motor learning adaptation[6–10]. These studies are critical for advancing rehabilitation strategies for individuals with motor function impairments, such as those caused by stroke or spinal cord injury[11–14]. Moreover, understanding motor learning adaptation may inform enhancements to artificial intelligence (AI) systems, as the brain adapts to disruptions more efficiently than current AI models[15–18].

In the past decade, brain-machine interface (BMI) paradigms have emerged as powerful tools for establishing causal links between neural activity and observable behaviors[19–22]. These paradigms enable researchers to investigate how neurons operate both individually and as part of populations to adapt neural activity during motor learning tasks[23–25]. In BMI tasks, individuals control external devices by volitionally modulating their neural activity. Since BMI learning establishes a causal mapping between neural signals (often from motor areas of the brain) and behavior, it can serve as a valuable model for studying motor learning adaptation. The controlled environment of BMI tasks simplifies the context of relevant motor variables to the neural signals and external actuator movements used in the BMI task, minimizing confounding factors and isolating the neural adaptation processes of interest[26]. Additionally, these tasks mimic natural motor learning through error-driven adjustments, providing insights into how the brain refines motor patterns in response to changes or perturbations in the external environment[27].

Researchers have found that population activity can often be characterized as having low-dimensional dynamics and that a pre-existing covariance structure constrains neural learning on short timescales[28]. This implies that learning to control existing population dynamics is more easily achieved compared to generating new activity patterns. In addition, other work suggests that we have a fixed repertoire of neural activity patterns, and that changes in population neural activity are induced by the reassociation of a selection of neural subspace latent variables[29]. Consequently, researchers have focused on understanding population-level neural dynamics, as population signals are more robust and better represent distributed cognitive and motor functions within local neuronal networks[30].

While these studies have highlighted the significance of population-level activity, understanding motor learning adaptation at the single-neuron level remains equally critical. Single neurons form the fundamental building blocks of neuronal networks and play a fundamental role in brain activity. Insights into single-neuron dynamics can reveal how individual neurons coordinate and respond to motor learning tasks, bridging the gap between population-level phenomena and individual neural mechanisms. For instance, while previous studies have investigated changes in firing rates at the single-neuron level during motor learning[31–38], questions about how individual neurons contribute to learning adaptation remain unanswered. Preferred direction changes in directionally-tuned single neurons have been observed[39,40], yet the coordination and contribution of individual neurons during motor learning adaptation remain poorly understood.

In this study, we address these gaps and provide additional granularity by examining how motor cortical neuron phenotypes explain the roles of individual neurons in BMI neurofeedback adaptation. Specifically, we categorize single neurons into narrow waveform (NW) neurons and broad waveform (BW) neurons. The classification based on waveform width is motivated by well-established electrophysiological properties of cortical neurons: narrow action potentials are thought to correspond to fast-spiking interneurons, which are predominantly inhibitory, while broad action potentials are associated with pyramidal neurons, which are primarily excitatory[41–45]. We demonstrate that NW neurons exhibit higher contributions to decoder output, greater coordination during task execution, and increased engagement during the learning process relative to BW neurons. By elucidating the distinct roles of NW and BW neurons, this study provides novel insights into the single-neuron mechanisms underlying neurofeedback adaptation, complementing existing knowledge of population-level neural dynamics.

## Results

We trained two rhesus macaques (monkeys A and B) to perform a two-dimensional BMI center-out task (Fig. 1A) which is a neurofeedback task. In this task, the animals controlled a cursor on a computer screen by volitionally modulating their neural activity, aiming to move the cursor to one of eight radially distributed peripheral targets to earn a juice reward. Neural activity was quantified as spike counts binned into small time intervals, which served as the high-dimensional input to the BMI decoder, while cursor velocity updates were the low-dimensional decoder output. A Kalman filter-based decoder, a widely used approach in BMI research, mapped the relationship between neural activity inputs and cursor velocity output dynamics[23,28,29,40,46].

**Figure 1.**
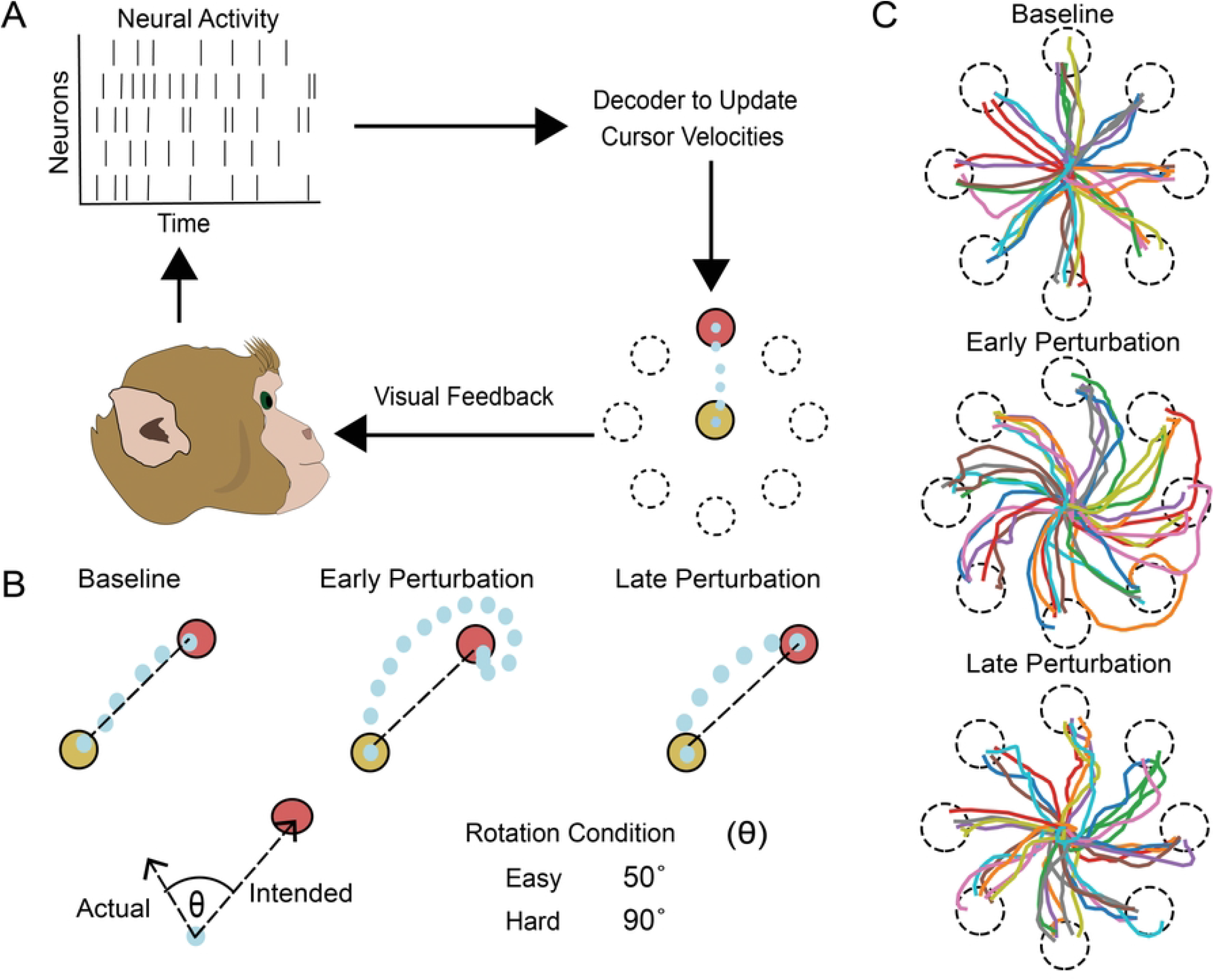
Overview of the BMI task and observed cursor trajectories. (A) Illustration of the BMI-based center-out task. Monkeys controlled a cursor on a screen by modulating neural activity based on visual feedback of resultant cursor movements. (B) The task was comprised of two blocks: baseline and perturbation. The decoder, which maps neural activity to cursor velocities, was rotated by 50 degrees or 90 degrees during the perturbation block. (C) Representative cursor trajectories to each of the possible target locations, depicted by the dashed line circles. In the baseline block, trajectories were relatively straight (top). During the early phase of the perturbation block (first 40 trials), trajectories were more curved, with longer distances from the center to the targets (middle). In the late phase of the perturbation block, trajectories became straighter again after learning adaptation process.

The BMI center-out task consisted of two distinct blocks: a baseline block and a perturbation block. Before the task, we trained the decoder using online data to establish an “intuitive mapping”[47,48]. This mapping aligned neural activity patterns with cursor dynamics in a way that mimicked natural motor control, enabling the monkeys to achieve proficient cursor movement control in the baseline block (see Methods). This block consisted of an equal number of trials per peripheral target. In the perturbation block, we introduced a rotation to the intuitive mapping by applying a rotation matrix to the Kalman gain in the BMI decoder (Fig. 1B), disrupting the established correspondence between neural activity and cursor movement. This perturbation mapping introduced a rotational mismatch, meaning the direction of cursor movement no longer aligned with the neural activity patterns that previously controlled it. This perturbation resulted in a misalignment between intended and actual cursor trajectories. The perturbation block comprised also consisted of an equal number of trials to the eight possible peripheral targets (Fig. 1B), requiring the monkeys to adjust their neural activity to compensate for the rotation for targets in all directions. These adjustments reflect motor learning and adaptation. We tested two rotation perturbation conditions: an “easy” condition with a 50-degree rotation and a “hard” condition with a 90-degree rotation.

As anticipated, both monkeys displayed fast and stereotypical target acquisition during the baseline block (Fig. 1C). In contrast, the early phase of the perturbation block was characterized by prolonged target acquisition times and curved cursor trajectories as the subjects struggled to compensate for the rotation (Fig. 1C). Over time, the monkeys adapted to the perturbation, as evidenced by straighter cursor trajectories in the late phase of the perturbation block.

### Neuron classification

Neural activity was recorded from the forelimb regions of the dorsal premotor cortex (PMd) and primary motor cortex (M1) using chronically implanted electrode arrays. Neurons with an average firing rate above 1 spike/s and trough-to-peak amplitude greater than 80 µV were selected as inputs to the BMI decoder. For the phenotype classification, we first measured trough-to-peak intervals for all these input neurons across sessions to quantify the typical width of their action potential waveforms (Monkey A: 1,195 neurons; Monkey B: 3,814 neurons). The distributions of trough-to-peak intervals were bimodal for both subjects, suggesting a mixture of activity generated from two categories of neurons based on action potential waveform width (Monkey A: p = 0.0274, Monkey B: p = 0.016; Hartigan dip test). Fitting a two-component Gaussian mixture model to spike-to-trough interval distributions significantly improved model fit compared to a single-component model (S1 Fig.).

Based on these results, neurons were categorized into three groups: narrow waveform (NW) neurons, broad waveform (BW) neurons, and unclassified neurons (Fig. 2A). Unclassified neurons had mid-range action potential waveform widths which could not confidently be categorized as narrow or broad and thus were excluded from further analysis. Categorization was determined by waveform trough-to-peak thresholds that were derived empirically from the respective distributions for each subject. In Monkey A, 27.2% (325/1195) were classified as NW neurons, 19.3% (737/3814) in Monkey B were identified as NW neurons. These proportions aligned with previous findings, which report NW neurons constituting 15% to 30% of the cortical neuron population[49–51]. The ratios of BW neurons in both monkeys were approximately 50% of total input neurons (42.93% in monkey A and 56.27% in monkey B).

**Figure 2.**
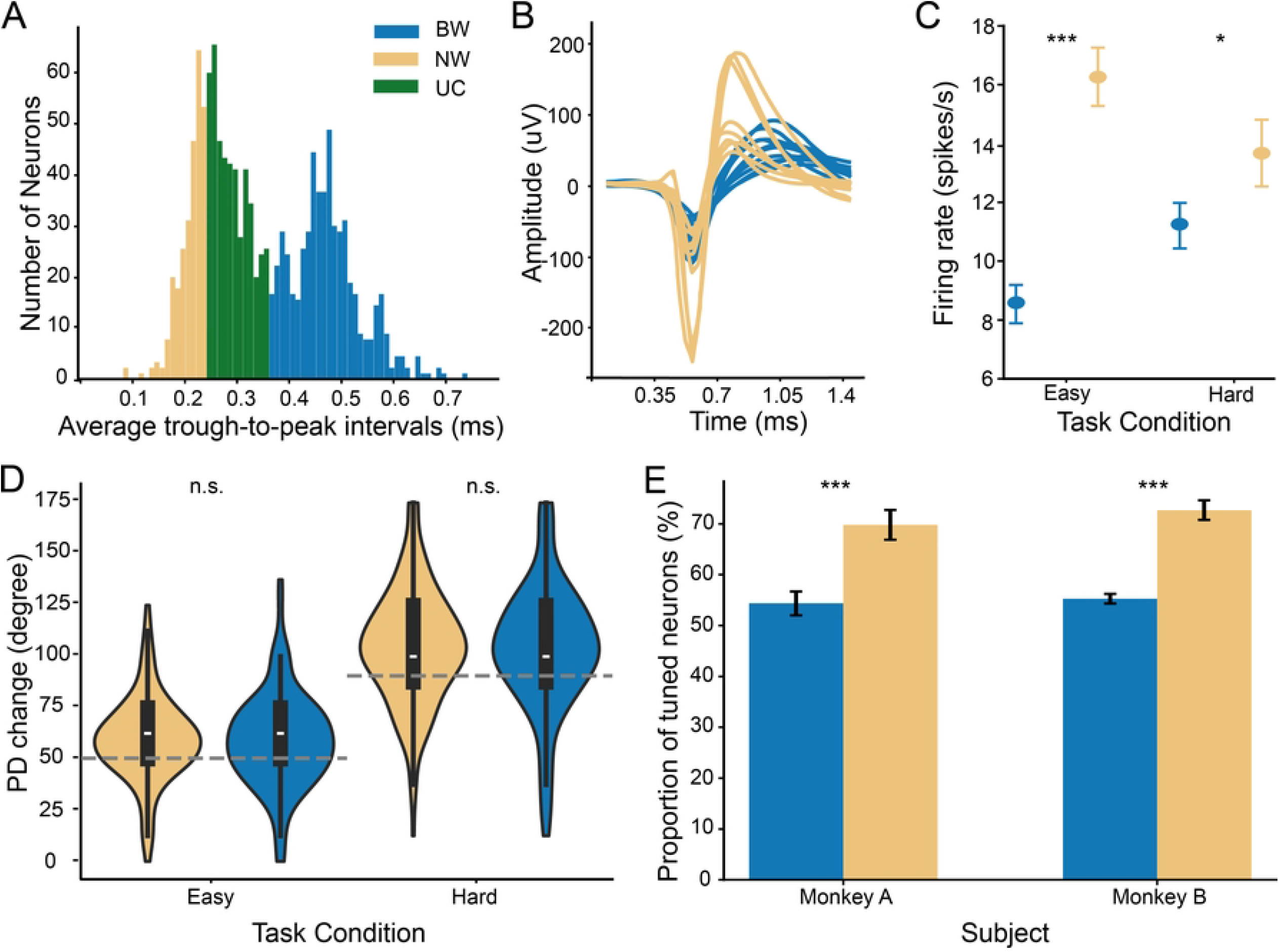
Neuron classification. (A) Distribution of trough-to-peak durations for three neuron groups: narrow waveform (NW, yellow), broad waveform (BW, blue), and unclassified (UC, green). The histogram represents all input neurons recorded in Monkey A. (B) Representative waveforms of NW (yellow) and BW (blue) neurons, aligned at the trough for comparison. (C) . Mean firing rates of NW and BW neurons during easy and hard task conditions. Orange markers indicate the mean firing rates of NW neurons, while blue markers represent those of BW neurons. Error bars denote the standard error of the mean (SEM). Statistical significance was observed (p = 9.4 × 10^−13^) for the easy condition; p = 0.031 for the hard condition). (D) Comparison of preferred direction (PD) changes between well-tuned neurons in NW and BW neurons. No significant differences in mean PD changes were observed between NW and BW neurons (easy condition: t = -0.211, p = 0.833; hard condition: t = -0.268, p = 0.789). However, the variance in PD changes increased significantly under the hard condition compared to the easy condition (Levene’s test: BW neurons, p = 0.002; NW neurons, p = 0.044). The grey dashed line indicated the rotation angle of the tasks. (E) Different proportions of well-tuned neurons in BW and NW neurons (monkey A: t=-4.106, p = 0.000113; monkey B: t=-8.129, p = 4.51*10^−12^), two-sample t-tests).

The mean firing rate of NW neurons during cursor movements was significantly higher than that of BW neurons (Fig. 2B) under both easy and hard task conditions (Monkey A: p <0.001 for the easy condition, p <0.05 for the hard condition; Monkey B: p<0.001 for both conditions; two-sided Wilcoxon rank-sum test). This high firing rate of NW neurons (Fig. 2C), which is typical of inhibitory neuron, is also consistent with findings from previous studies[50,52,53].

Previous studies have shown that neurons change their preferred direction (PD) during visuomotor rotation tasks, suggesting that neurons adapt in response to altered movement mappings[38,39]. Despite evidence of PD shifts during visuomotor adaptation, whether distinct neuronal phenotypes exhibit these changes in a uniform or differential manner remains unresolved. We calculated the PD of NW and BW neurons separately for the baseline and perturbation blocks by fitting tuning curves[39] (see Methods). Neurons were well-tuned and included in the PD analysis if their tuning curve fits in both blocks had *R*^2^ values greater than 0.5. More than 60% of NW and BW neurons were well-tuned in both blocks (S2 Fig.), and this proportion remained consistent across task difficulty levels, with minor differences likely attributable to session-to-session variations in neuron recordings.

Both NW and BW neurons demonstrated mean PD changes corresponding to the rotation angle (50° for the easy condition and 90° for the hard condition; Fig. 2D). There was no significant difference in median PD changes between NW and BW neurons under either task condition (p > 0.05, two-sample t-test). However, the variance in PD changes increased significantly under the hard task conditions for both NW and BW neurons (p < 0.01 for BW neurons; p < 0.05 for NW neurons; Levene’s test). The similar mean PD changes in well-tuned neurons for both types of neurons suggested that these well-tuned neurons were involved in comparable underlying adaptation processes. However, the proportion of well-tuned neurons in the NW and BW neuron subpopulations were significantly different in both subjects (Fig. 2E). In both subjects, NW neurons had a significantly higher proportion of well-tuned neurons than BW neurons (p < 0.001 for both subjects). While well-tuned neurons in both phenotypes shifted their PD in alignment with the rotation angle in the task, their different proportions suggested that NW and BW neurons may engage distinct mechanisms during motor learning.

### Unit contribution

To investigate how the physiological properties of neurons influence their roles in the behavioral task, we evaluated the contributions of BW and NW neurons to neurofeedback control. We quantified these contributions by defining the unit contribution (*C*_*i*_) as below:

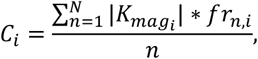

where 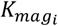 represents the magnitude of the Kalman gain:

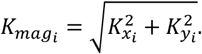

Here, *C*_*i*_ is the unit contribution for neuron *i* averaged across *N* trials; 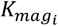 is the magnitude of Kalman gain (see Methods) associated to neuron *i*; and *fr*_*n,i*_ is mean firing rate of neuron *i* on trial *n*. Given that the intuitive decoder was trained without incorporating neuron phenotype, differences in the Kalman gain magnitude (*K*_*mag*_) between NW and BW neurons were not expected. Indeed, our analysis confirmed that the distributions of *K*_*mag*_ did not significantly differ between the two neuronal populations in either monkey (Fig. 3A, B; p > 0.05 for both), suggesting that the gain magnitude was independent of neuron type.

**Figure 3.**
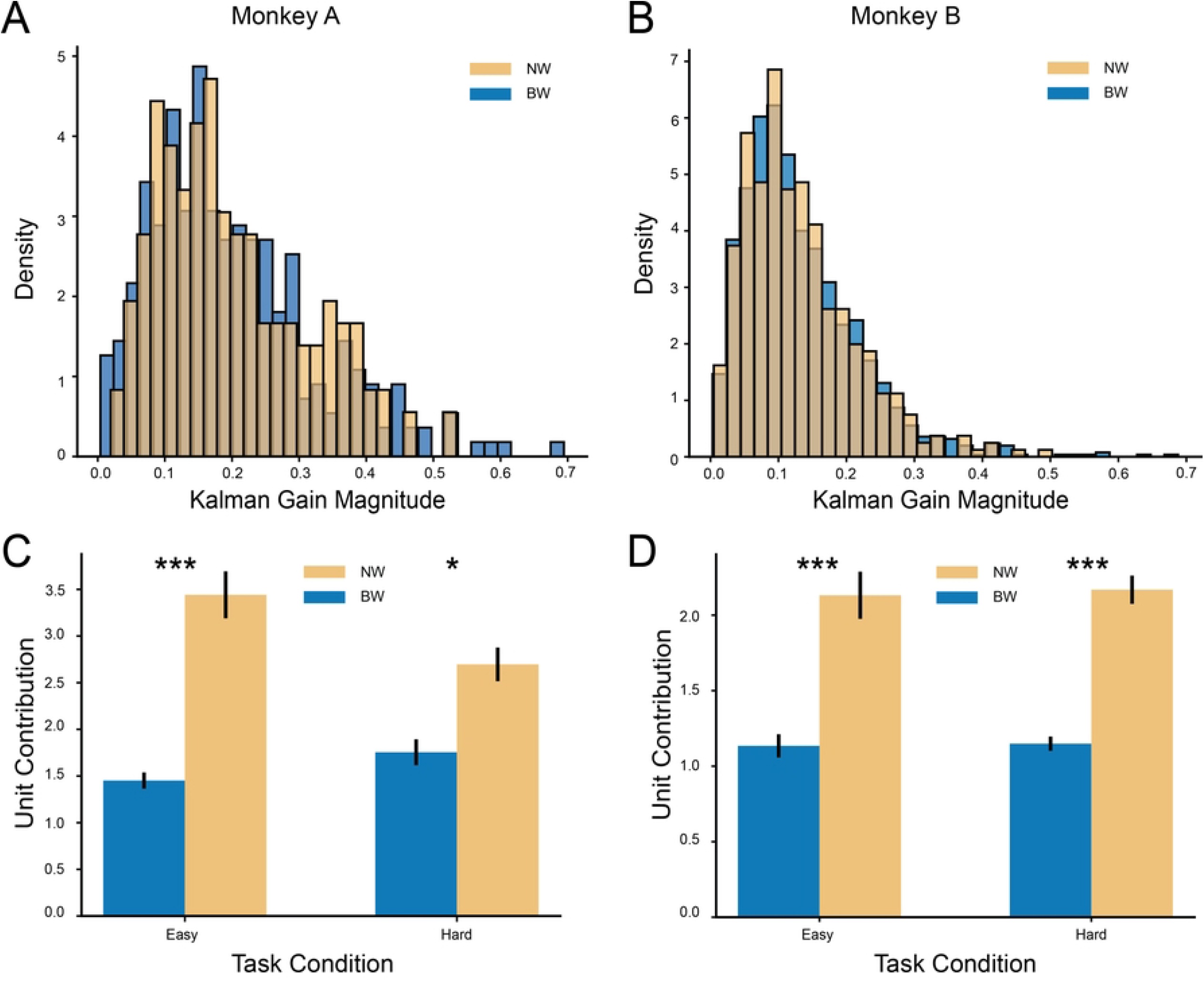
NW neurons exhibit higher unit contributions compared to BW Neurons. (A), (B) Distributions of Kalman Gain magnitudes for narrow waveform (NW) and broad waveform (BW) neurons in Monkey A and Monkey B. No significant differences were observed in the Kalman Gain magnitudes between NW and BW neurons in either monkey (Monkey A: t = 0.078, p = 0.938; Monkey B: t = 0.339, p = 0.735; two-sample t-tests). (C), (D) Comparisons of average unit contributions from NW and BW neurons in Monkey A and Monkey B under easy and hard task conditions. In both conditions, NW neurons exhibited significantly higher contributions than BW neurons. For Monkey A, NW neurons showed greater contributions in the easy condition (z = 10,215, p = 4.099 × 10e-10) and the hard condition (z = 22,169, p = 0.015). For Monkey B, NW neurons also had significantly higher contributions in the easy condition (z = 100,963, p = 2.816 ×10^−30^) and the hard condition (z = 157,074, p = 3.121 ×30; Mann-Whitney U tests).

Despite the similarity in *K*_*mag*_, we found that NW neurons consistently exhibited greater unit contributions than BW neurons across both task difficulty conditions (Fig. 3C, D; Monkey A: p < 0.001 in easy condition, p<0.05 in hard condition, Monkey B: p < 0.001 in both task conditions). These findings suggest that NW neurons play a dominant role in neurofeedback control, which can be attributed to their higher firing rates that amplify their influence within the decoder.

### Coordination in BW and NW neurons at population level

Neuron contributions revealed the unique roles of NW and BW neurons in shaping decoder output. However, the mechanisms underlying their adaptation during motor learning remain unclear. Previous research suggests that neural activity during motor adaptation at the population level can be represented in a low-dimensional space[28,29]. Moreover, researchers have shown that motor learning within the same low-dimensional space is induced by reassociation of latent variables[29]. Moreover, previous research indicates that as performance improves, neurons tend to exhibit more synchronized firing patterns[23,54]. These synchronized patterns may reflect the consolidation of neural population dynamics, which can be examined through the variance of neural activity within low-dimensional subspaces. Despite this, how different neuron types coordinate during motor learning adaptation has not been explored.

To measure neural coordination, factor analysis (FA) was performed on population neural activity to separate variance into two components: shared variance and private variance[23]. Shared variance reflects correlated firing patterns across the population, while private variance represents variability unique to each neuron[55,56]. The ratio of shared variance over total variance (SOT) is treated as a measurement to quantify the relative amount of coordinated neural activity[23,56]. To evaluate how BW and NW neurons adapt during motor learning, we calculated the partial shared-over-total variance (pSOT) of each phenotypical neuron group (see Methods) since the pSOT ratio assesses the extent to which each subpopulation contributes to the overall change in coordinated activity.

Under both easy and hard task conditions, NW neurons consistently demonstrated significantly higher pSOT ratios compared to BW neurons (three-way ANOVA, p < 0.001 for both monkeys; Fig. 4). This indicates that NW neurons demonstrated greater coordination within their subpopulation than BW neurons. Additionally, in Monkey B, pSOT ratios were significantly higher under hard conditions compared to easy conditions (Tukey HSD test, p < 0.05), suggesting that task difficulty may influence neural coordination. Notably, no significant differences in pSOT were observed between baseline and perturbation blocks for either NW or BW neurons. Furthermore, the latent structure of neural activity, as measured by the low-dimensional repertoire, did not differ significantly between baseline and perturbation blocks (S3 Fig.). These findings demonstrate that NW neurons play a crucial role in motor adaptation by exhibiting stronger coordination, especially under challenging task conditions.

**Figure 4.**
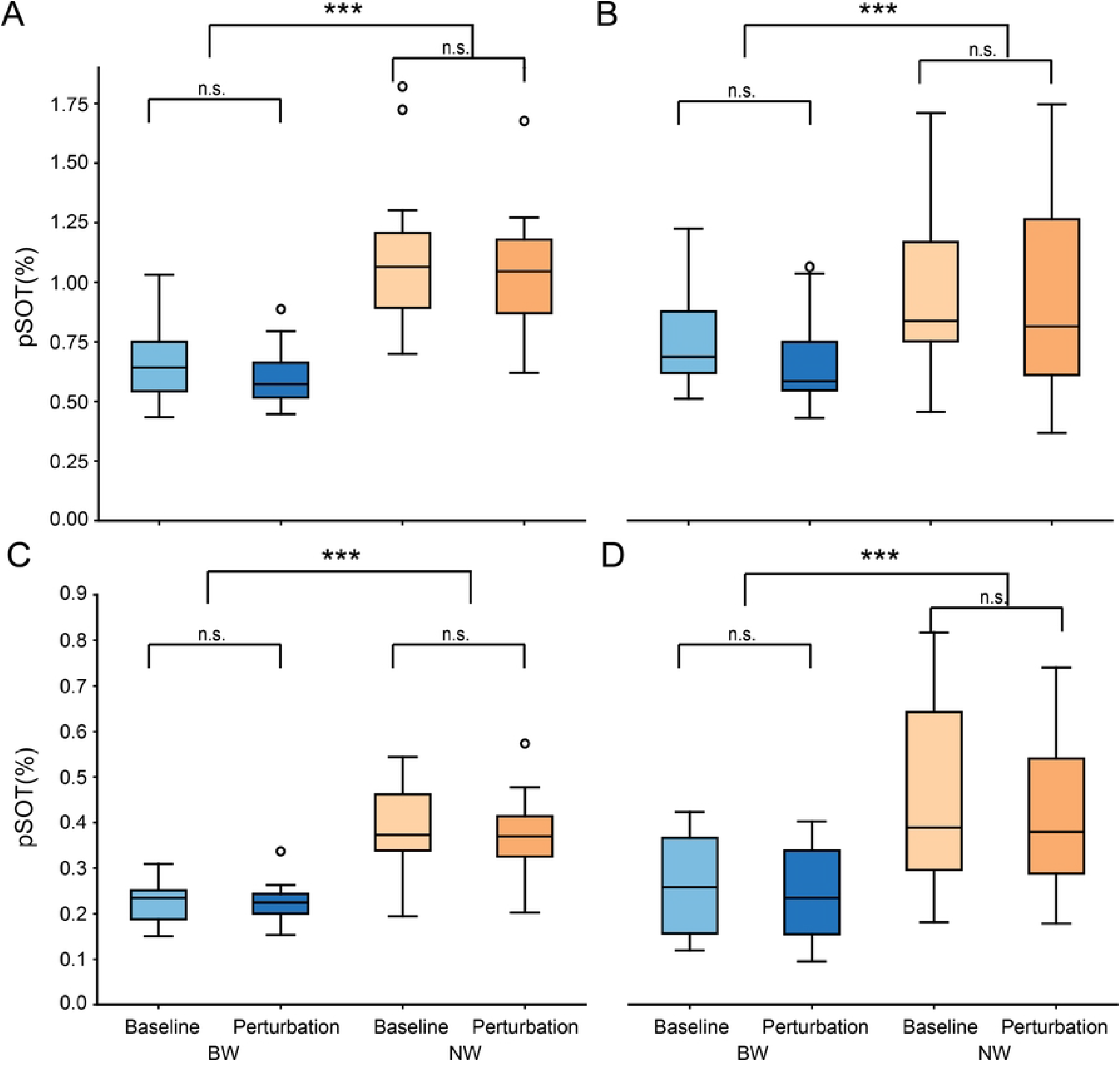
NW neurons show higher coordination than BW neurons through higher pSOT. (A), (B) These were pSOT results for monkey A. (A) was under easy condition while (B) was under hard condition. Three-way ANOVA of Neuron type (BW and NW), task blocks (baseline and perturbation) and task condition (easy and hard) was applied to Monkey A. NW neurons had significant higher pSOT values than BW neurons (p = 3.13 ∗ 10^−8^). No significant differences in pSOT were observed between the baseline and perturbation blocks (F= p = 0.067). Additionally, in Monkey A, there was no significant difference in pSOT between the easy and hard task conditions (p = 0.108). Same three-way ANOVA was applied to Monkey B data. (C) was Monkey B under easy task condition while (D) was under hard task condition. Significant difference was shown between neuron type (p = 2.28 ∗ 10^−13^ for Monkey B) while there was no difference between task blocks (Monkey B: p = 0.209). But a significant difference was detected between easy and hard task condition in monkey B (p = 0.012).

### Differential Responses of BW and NW Neurons at the Single-Neuron Level

While FA provided insights into the coordination of neural subpopulations, the trial-by-trial neural responses at the single-neuron level remained unclear. Previous studies have demonstrated that the percentage of explained variance (*ω*^2^) is an effective measure of neural coding changes under varying task conditions[49]. Since FA did not identify differences in neural activity between easy and hard task conditions in Monkey A, we applied *ω*^2^ to compare single-neuron responses during trials across neuron types. Neural recordings revealed substantial variability in firing rates across neurons, depending on cursor movement direction. To account for this, *ω*^2^ was calculated separately for all eight target locations.

To minimize variability in *ω*^2^ calculations, we analyzed the last 20 trials of the baseline block and the first 20 trials of the perturbation block for each target location. This comparison assessed the sensitivity of neural activity changes to motor learning adaptation in response to the perturbation. High *ω*^2^ values indicated stronger neural responsiveness during the learning adaptation period. Both BW and NW neurons exhibited significantly stronger encoding of cursor movements during the learning period compared to the center-hold period (Fig. 5A). Within trials, *ω*^2^ values for both neuron types showed an initial rapid increase followed by a slower decay in both easy and hard conditions. Additionally, NW neurons demonstrated higher average *ω*^2^ values than BW neurons across all target locations and task conditions in both subjects (Fig. 5B, 5C; p < 0.001, two-sample t-test in both monkey A and B), suggesting that NW neurons play a more prominent role in representing motor adaptation information. Moreover, both NW and BW neurons had significant higher *ω*^2^ values in hard condition than in easy condition (p < 0.001, two-sample t-test in both monkey A and monkey B). This indicated that when the task was more challenging, both neurons were more responsive during the task.

**Figure 5.**
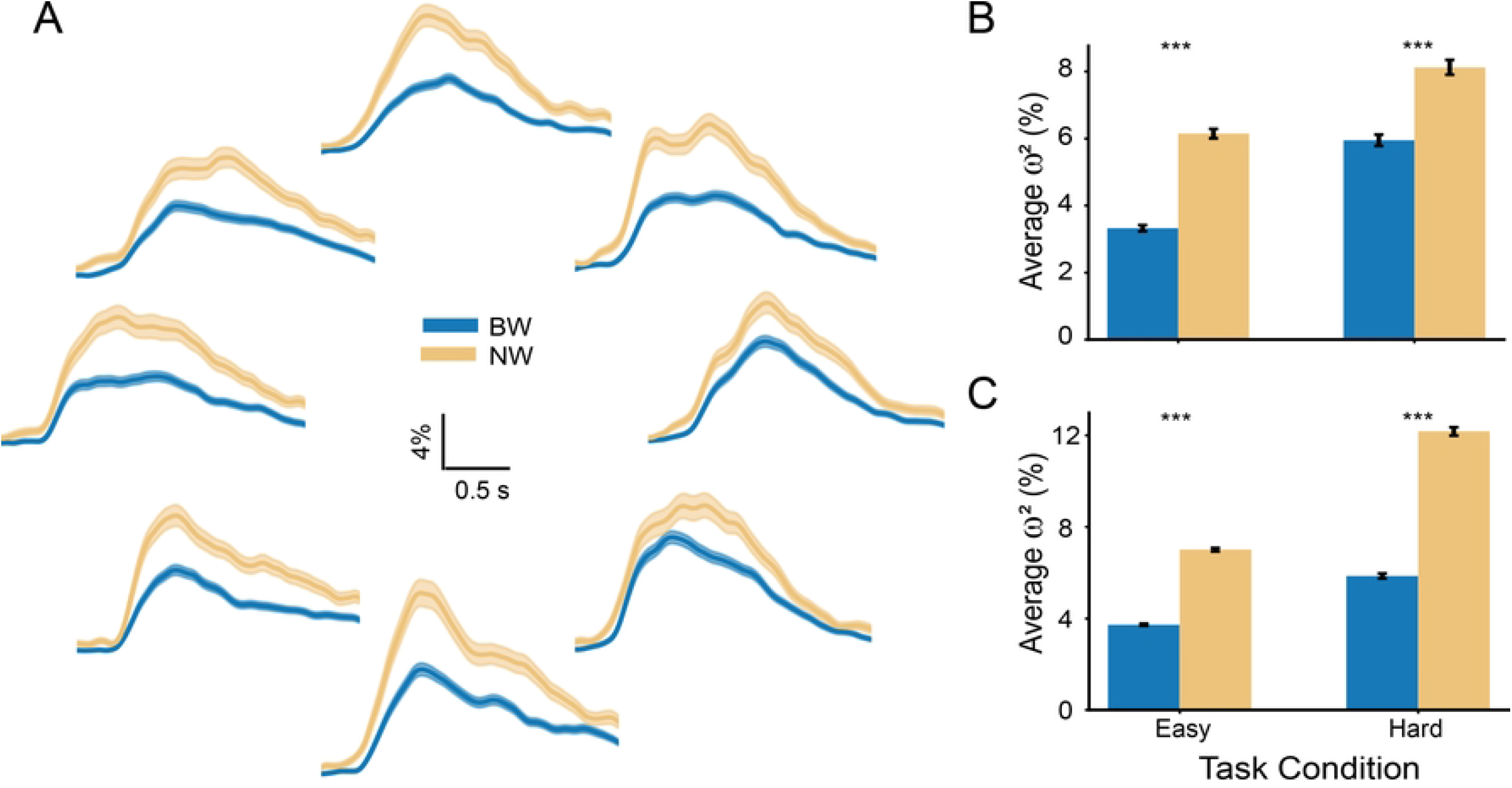
NW neurons had higher *ω*^2^ than BW neurons during cursor moving period. (A) *ω*^2^ values trends within trial time between BW (blue) and NW (yellow) neurons. The subplots’ locations represent the corresponding target location on the screen. Cursor movement period was from “Hold” state to “Target” state. Here, we restricted the visualization for trial time from 0.2s before “Hold” state and 2s after “Hold” state. (B), (C). NW (yellow) neurons have significantly higher average *ω*^2^ values than BW (blue) neurons in both task conditions. (monkey A: t =-14.655, p = 5.103* 10^−47^ in easy condition, t = -7.938, p = 2.745*10^−15^ in hard condition; monkey B: t = -20.153, p = 1.546*10^−88^ in easy condition, t = -31.005, p = 1.268*10^−203^).

We found that both NW neurons reached their peak *ω*^2^ values faster than BW neurons in both subjects. Moreover, both types of neurons show faster response in hard task condition compared to the easy task condition (Fig. 6A, B; monkey A: neuron types: p < 0.05; task condition: p < 0.001. Monkey B: neuron types: p << 0.001; task condition: p << 0.001, two-way ANOVA). Also, NW neurons shown greater *ω*^2^ values during the learning period compared to BW neurons. Furthermore, both neurons increased maximum *ω*^2^ values significantly during more challenging task conditions (Fig 6C,D. monkey A: neuron types: p << 0.001; task conditions: p << 0.001; monkey B: neuron types: p << 0.001; task conditions: p << 0.001,two-way ANOVA). These results suggest that the observed variance in firing rates between the last 20 trials of the baseline block and the first 20 trials of the perturbation block was driven by the induced rotation in the task. In other words, changes in firing rates for both neuron types are likely attributable to motor learning adaptation. Overall, *ω*^2^ served as a valuable metric for quantifying neural responses during motor learning adaptation. NW neurons were found to be more responsive and engaged during the learning period compared to BW neurons, and both neuron types demonstrated increased responsiveness during more challenging tasks.

**Figure 6.**
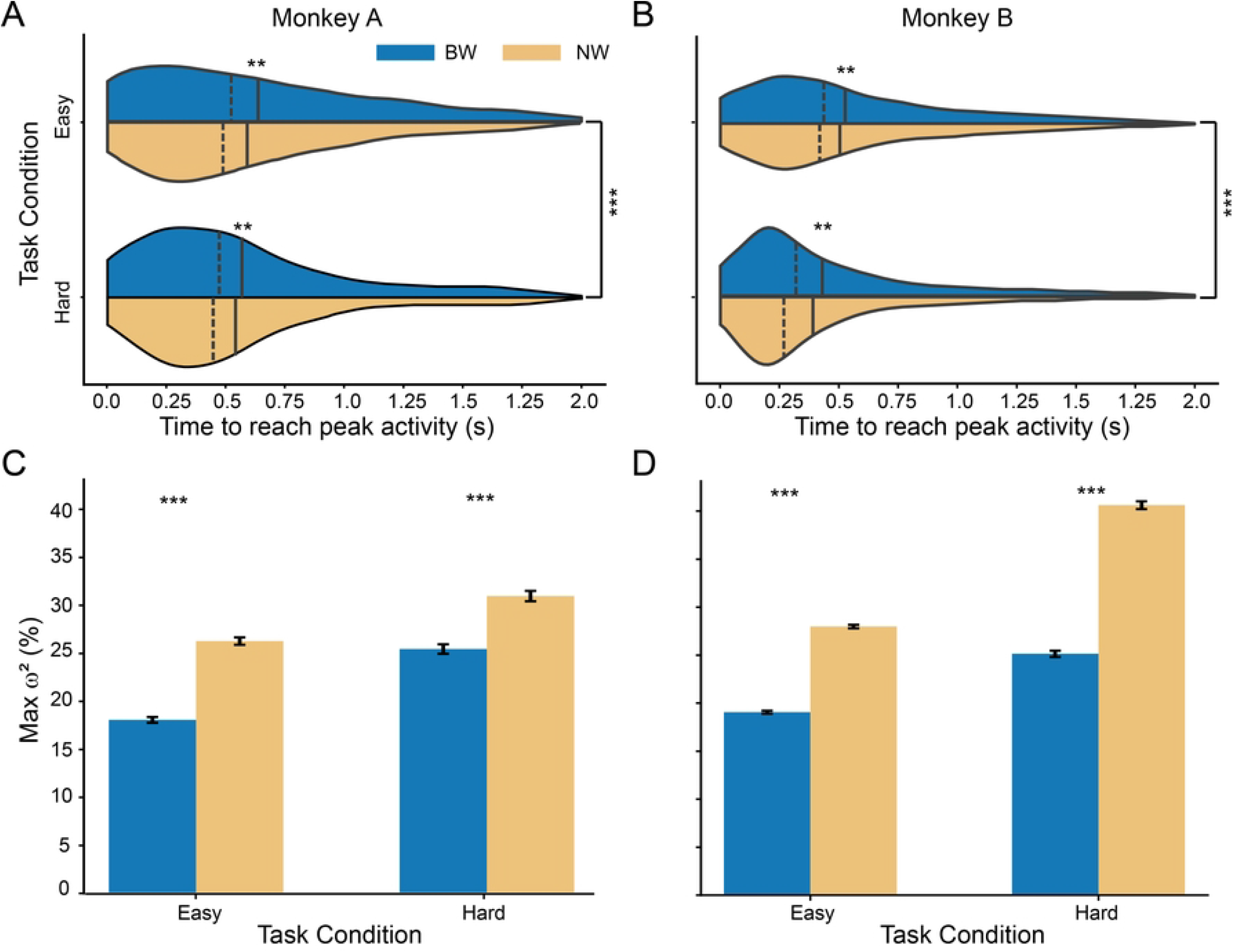
Neuron phenotype and task difficulty affected neurons’ response. (A), (B) Time to reach peak *ω*^2 values were different between NW (yellow) and BW (blue) neurons (monkey A: neuron type: F=6.401, p =0.011 task difficulty: F=23.650, p=10^−5^; monkey B: neuron type: F=28.178, p = 1.117 ∗ 10^−7^ task difficulty: F=410.246, p=0, two-way ANOVA test).). (C), (D) Max *ω*^2^ values comparison between NW and BW neurons(monkey A: neuron type: F=189.91, p = 1.31 ∗ 10^−42^; task difficulty: F=291.34, p=6.13 ∗ 10^−64^; monkey B: neuron type: F=1278.34, p = 2.12 ∗ 10^−272^ task difficulty: F=2125.36, p=0, two-way ANOVA test).).

## Discussion

In this study, we investigated how individual neurons participate in motor learning adaptation, focusing on whether these neurons exhibit uniform responses during the adaptation period within a precisely defined BMI task. Neurons were classified into two groups, NW neurons and BW neurons, based on their waveform shapes (trough-to-peak intervals; Fig. 2). Our results demonstrate that NW neurons contribute more to the decoder output (Fig. 3) and exhibit greater pSOT indicative of higher coordination during task execution (Fig. 4). NW neurons also display higher *ω*^2^ values, reflecting greater engagement during the learning period. Moreover, larger rotations in the intuitive mapping elicited more substantial changes in neural activity in both neuron groups compared to smaller rotations.

Our classification of neurons into NW and BW groups based on their trough-to-peak intervals aligns with previous studies, which have identified a bimodal distribution of spike waveform widths indicative of the separation between excitatory cells and inhibitory interneurons in the cortex[49–51,57]. Specifically, NW neurons, which exhibit higher firing rates than BW neurons, are consistent with prior observations[49,58]. These characteristics support the hypothesis that NW neurons are likely inhibitory interneurons. Moreover, approximately 70% of the NW neurons in the dorsal premotor cortex (PMd) and primary motor cortex (M1) exhibited preferred directions, with many adjusting their preferred directions following task rotation. This proposition is further corroborated by findings in the primary visual cortex showing that interneurons also have preferred orientations[59–61]. In this study, inhibitory neurons exhibited directional tuning, which may be caused by the inputs from the pyramidal neurons through the recurrent cortical circuitry. M1 contains extensive recurrent excitatory-inhibitory loops[62,63]. Pyramidal neurons excite inhibitory neurons, which in turn regulate pyramidal neuron activity. This recurrent architecture means that the directional tuning seen in inhibitory neurons is likely driven by the activity of the pyramidal neurons rather than independent computation.

Our findings revealed differences in coordination between NW and BW neurons during motor learning adaptation. NW neurons exhibited higher pSOT ratios, indicating greater coordination within their population. Furthermore, we observed no significant changes in the latent variable repertoire between the baseline and perturbation blocks, consistent with previous findings[29]. This stability in the latent variable repertoire likely reflects changes in the preferred direction of individual neurons without significant alterations in local neuronal networks. These results underscore the adaptability of individual neurons in response to task perturbations, while broader network structures remain relatively stable.

There are limitations to this study that should be noted. Firstly, we did not record neural activity from upstream brain regions, which could provide direct insights into the extent of changes that are attributable to upstream inputs. M1 and PMd receive projections from multiple upstream areas, including the somatosensory cortex, supplementary motor area, and thalamus[64,65]. Without direct measurements from these regions, it remains unclear to what extent the observed neural changes are attributable to modifications in upstream inputs versus intrinsic plasticity within local cortical networks. Second, the classification of unclassified neurons remains an open issue, as they account for approximately one-third of all input neurons in the decoder. To ensure accurate classification, we prioritized precision over quantity for the NW neuron type, which led to a higher proportion of unclassified neurons. Notably, previous studies have reported that pyramidal cells constitute 70–85% of all neurons in the mammalian cortex[66,67]. However, in our dataset, BW neurons comprise only about 50% of the total recorded neurons. This discrepancy arises because NW neurons exhibited a much narrower distribution compared to BW neurons. This narrower distribution of NW neurons forces stricter classification criteria, leading to more BW neurons being labeled as unclassified neurons to avoid misclassification. In this study, we relied solely on electrophysiological recordings and lacked additional methods to further differentiate these neuronal types. Future studies incorporating complementary recording techniques may provide greater resolution in distinguishing these two neuronal populations.

Our findings emphasize the differential contributions of NW and BW neurons during BMI motor learning adaptation. Both neuron groups contain well-tuned neurons capable of adjusting their tuning properties in response to varying rotation perturbations, but NW neurons exhibit higher coordination and engagement during motor learning. Additionally, both groups undergo greater changes in neural coordination as task complexity increases. Understanding the distinct roles of different neuronal groups during neural adaptation provides valuable insights into the mechanisms underlying rapid learning. These findings may guide the development of more sophisticated brain-inspired artificial networks and contribute to achieving more stable and precise BMI control systems.

## Method

### Experimental models and procedures

Two male rhesus macaques (Macaca mulatta, A: 6 years old, 9.5 kg, B: 6 years old, 8.8 kg) participated in the experiments. All procedures performed were approved by the Institutional Animal Care and Use Committee (IACUC) of The University of Texas at Austin. Both subjects were implanted with chronic arrays (Innovative Neurophysiology, Inc., Durham, NC) in their left hemisphere primary motor (M1) / pre-motor (PMd) area of cortex (Monkey A: 64-channel chronic array; Monkey B: 128-channel chronic array). Prior to the surgery, we used structural MR images to determine the targeted implantation area.

### Neural recordings

Neural activity was recorded by Grapevine Neural Interface Processor (Ripple Neuro, Salt Lake City, UT). Spike activity was recorded with a sampling rate of 30 kHz and we included only units with a firing rate greater than 1 spike/s in our BMI decoders. On average 37.6 ± 6.2 units in monkey A and 91.6 ± 19.3 units in monkey B were used. Spikes were binned into 100 ms time windows to generate spike count vectors for decoder inputs and the BMI decoder was output updated at a rate of 10 Hz.

### Behavioral tasks

Subjects were trained to perform a 2-D center-out BMI task beginning with holding the cursor (circle, radius 5 mm) at the center target (circle, radius 20 mm) for 200 ms. One of eight peripheral targets which are uniformly distributed around a circle (radium 100 mm) is then shown on screen to cue the cursor destination. The animal volitionally controls their neural activity to move the cursor from the previous holding center position to this shown target. After successfully holding the cursor at the peripheral target for 500 ms, a liquid reward is delivered to the subject’s mouth. If the subject cannot complete this within 10 s, the trial is considered to be unsuccessful. Each experiment session starts with a 3-minute visual feedback (VFB) task. During the VFB task, the subject passively watches a computer-controlled simulation of the BMI task while receiving the juice reward for all completed trials. We use neural activity recorded during VFB to determine the units used for the BMI decoder in the brain-controlled neurofeedback task. After this VFB task, the subject performs a 10-minute center-out BMI task with gradually decaying computer assistance in order to optimize the decoder parameters. When this 10-minute task ends, the intuitive mapping decoder is trained well, and the subject starts the main center-out BMI task. In the main task, there are two blocks: baseline and perturbation. The baseline block contains 336 trials (42 trials per peripheral target), and the perturbation block contains 384 trials (48 trials per peripheral target). In the baseline block, the decoder used is the well-trained intuitive mapping decoder. In the perturbation block, a defined rotation matrix is applied to the intuitive mapping decoder to impose a visuomotor rotation on the cursor. To gain the liquid juice reward, the animals must adapt their neural activity to compensate for the perturbed output from the decoder.

### Kalman filter decoder

We use a Kalman filter as the decoder[23,40,54] to map neural activities and cursor velocities, shown as below:

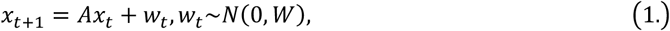

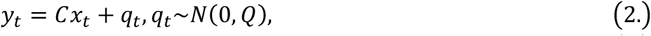

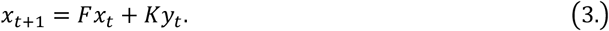

The cursor state *x*_*t*_ ∈ *R*^5^ is a 5-by-1 vector [*pos*_*x*_,*pos*_*y*_,*vel*_*x*_,*vel*_*y*_,1]^*T*^ and *A* is a 5-by-5 matrix, capturing the physics of cursor position and velocity. The observed neural activity *y*_*t*_ ∈ *R*^*N*^, where N is the number of input neuron units. *C* represents the relationship between neural activity and cursor movement. Gaussian noise is included as *w*_*t*_and *q*_*t*_. In equation (3), *K*∈ *R*^5×*N*^ is a 5-by-N Kalman Gain matrix. Since we use the neural activity to decode the cursor velocity, the cursor position is updated by the velocity multiplied by the time interval. The decoder is updated at a rate of 10 Hz, so the time interval used equates to 100 ms.

In the perturbation block, a rotation matrix is applied to *K* to rotate the intuitive mapping decoder, which is shown as below:

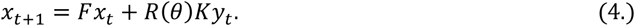

The rotation angle is *θ*, which was set to be either 50° or 90°.

### Neural waveform classification

To categorize the cells in our recordings into broad and narrow waveform spiking types, we employed a method established in previous research[53,68,69]. Each cell’s average waveform was subjected to cubic interpolation, enhancing the sampling resolution from 33 μs to 1.65 μs. The interpolated waveforms were then normalized and aligned to the primary trough. The trough-to-peak duration, defined as the interval between the main trough and the subsequent waveform peak, served as the classification metric.

To statistically evaluate the bimodality of the trough-to-peak distribution, we applied two variations of the Hartigan dip test: the original version and a calibrated variant with enhanced sensitivity. Additionally, we used Akaike’s and Bayesian information criteria to compare whether the distribution was better represented by a one- or two-Gaussian model. For the two-Gaussian model, we defined two cutoffs to segment the distribution into three regions. The first cutoff corresponded to the point where the likelihood of classification as a narrow cell was at least 10 times higher than as a broad cell[49,53]. The second cutoff was set where the probability of classification as broad exceeded that of narrow by the same factor. Cells within the region between these cutoffs were deemed unclassifiable. Classification cutoffs for NW and BW neurons were consistent across both monkeys (Supplementary Fig. S1; cutoff for NW neurons: 0.243 ms, cutoff for BW neurons: 0.36ms in monkey A; cutoff for NW neurons: 0.24ms, cutoff for BW neurons: 0.373ms in monkey B), suggesting that these classifications represent distinct neuronal phenotypes across subjects rather than artifacts of feature selection.

### Tunning curves

We use a tuning curve to fit the neural responses of different moving directions[39]. The tuning curve is defined as a cosine function, showing below:

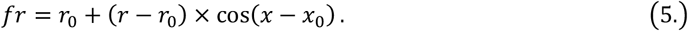

Preferred direction is the movement direction with highest firing rate and modulation depth is the difference in firing rate between peak and trough of the fitted tunning curve.

### Factor analysis

Factor analysis (FA)[23] decomposes signals into shared and private components. We apply FA to the collected neural data to obtain shared variance to total variance and partial percent shared variance to total variance (PSOT), which are defined as:

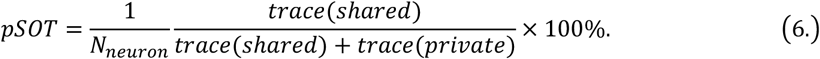

*pSOT* is the mean percent shared variance per unit. After cross-validation, we determined that 6 factors for both monkey A and B were appropriate.

### Percentage of explained variance

We used the percentage of explained variance (*ω*^2^) to measure the information provided by NW and BW neurons. This method enabled us to determine the proportion of variance in the neurons’ firing rates attributed to the tested variable[49,70,71]. *ω*^2^ is defined as:

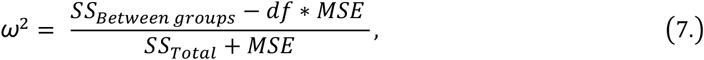

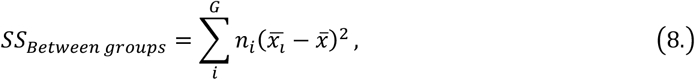

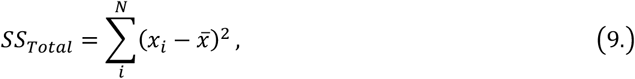

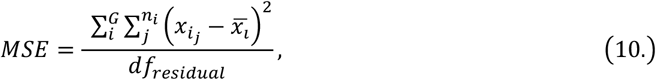

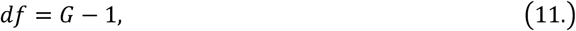

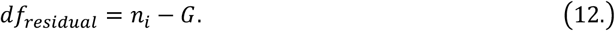

*G* is the total number of groups, and N represents the total number of trials for the cells. *af* denotes the degrees of freedom.

## Supporting information

**S1 Fig. Two mixed Gaussian models of trough-to-peak intervals**. The reductions in the Akaike Information Criterion (Monkey A: -1592 to -1860; Monkey B: -5265 to -5594) and the Bayesian Information Criterion (Monkey A: - 1581 to -1835; Monkey B: -5253 to -5563) show that two mixed Gaussian models were better than a single Gaussian model.

**S2 Fig. Pie plots showing ∼60% of all neurons are both tuned neurons**. In both task conditions, around 60% of total neurons are tuned in both baseline and perturbation blocks (yellow). Only around 10% of total neurons were only baseline tuned (red) or perturbation tuned (purple). Around 15% of total neurons were not sensitive to directions in both baseline and perturbation.

**S3 Fig. Example latent repertoire**. There was no significant difference between the latent repertoire between baseline and perturbation (t2=1.75e-29, p=1, Hotelling’s T^2 test)

## Data availability statement

This information will only be available after acceptance.

